# Antibody recognition of CD4-induced open HIV-1 Env trimers

**DOI:** 10.1101/2022.07.27.501785

**Authors:** Zhi Yang, Kim-Marie A. Dam, Jonathan M. Gershoni, Susan Zolla-Pazner, Pamela J. Bjorkman

**Author notes:** Zhi Yang and Kim-Marie A. Dam contributed equally to this work. Author order was determined by the two co-first authors after negotiation. Address correspondence to Pamela J. Bjorkman.

## Abstract

HIV-1 envelope (Env), a heterotrimer of gp120-gp41 subunits, mediates fusion of the viral and host cell membranes after interactions with the host receptor CD4 and a coreceptor. CD4 binding induces rearrangements in Env trimer, resulting in a CD4-induced (CD4i) open Env conformation. Structural studies of antibodies isolated from infected donors have defined antibody-Env interactions, with one class of antibodies specifically recognizing the CD4i open Env conformation. Here, we characterize a group of monoclonal antibodies isolated from HIV-1 infected donors (V2i mAbs) that display characteristics of CD4i antibodies. Binding experiments demonstrate that the V2i mAbs preferentially recognize CD4-bound open Env trimers. Structural characterizations of V2i mAb-Env-CD4 trimer complexes using single-particle cryo-electron microscopy show recognition by V2i mAbs using different angles of approach to the gp120 V1V2 domain and the β2/β3 strands on a CD4i open conformation Env with no direct interactions of the mAbs with CD4. We also characterize CG10, a CD4i antibody that was raised in mice immunized with a gp120-CD4 complex, complexed with Env trimer and CD4. CG10 exhibits similar characteristics to the V2i antibodies: i.e., recognition of the open Env conformation, but shows direct contacts to both CD4 and gp120. Structural comparisons of these and previously characterized CD4i antibody interactions with Env provide a suggested mechanism for how these antibodies are elicited during HIV-1 infection.

**Importance:** The RV144 HIV-1 clinical vaccination trial showed mild protection against viral infection. Antibody responses to the V1V2 region of HIV-1 Env gp120 were correlated inversely with the risk of infection. In addition, antibodies targeting V1V2 have been correlated with protections from SIV and SHIV infections in non-human primates. We structurally characterized V2i antibodies directed against V1V2 isolated from HIV-1 infected humans in complex with open Env trimers bound to the host receptor CD4. We also characterized a CD4i antibody that interacts with CD4 as well as the gp120 subunit of an open Env trimer. Our study suggests how V2i and CD4i antibodies were elicited during HIV-1 infection.

## Introduction

The HIV-1 envelope glycoprotein (Env), the only viral protein on the surface of the virus, interacts with target cells to mediate fusion between the viral and host cell membranes, a process that marks the initiation of HIV-1 infection (1–3). The trimeric Env is composed of three copies of gp120-gp41 heterodimers (1). To infect cells, the gp120 subunit interacts with the host cell receptor CD4 and undergoes a series of conformational changes that lead to the exposure of the binding site for a host cell coreceptor, either CCR5 or CXCR4 (4, 5). Env binding to its coreceptor results in further changes including the insertion of the gp41 N-terminal fusion peptide into the host cell membrane and subsequent fusion of the viral and host cell membranes (1).

Soluble versions of HIV-1 Env ectodomains that were stabilized in a closed, prefusion conformation (SOSIP.664 trimers) (6) have been used to characterize interactions between antibodies and Env in the presence and absence of soluble CD4 (sCD4) (7), with the clade A BG505 SOSIP (6) and the clade B B41 SOSIP (8) being commonly used in structural studies. Structural characterizations using X-ray crystallography or single-particle cryo-electron microscopy (cryo-EM) depicted different conformational states (Fig. 1A) including: (i) a closed, pre-fusion state in which the gp120 V1V2 region is positioned at the trimer apex, shielding the V3 loop and the coreceptor binding site (7, 9–11), (ii) an “occluded open” state in which the trimer is open due to outward rotation of the gp120 protomers, but with no local structural rearrangements in the V1V2 and V3 regions with respect to the remainder of the gp120 subunit (12, 13), and (iii) a CD4-induced, fully-open state in which the gp120 protomers rotate outwards with the V1V2 region displaced by ~40 Å to the sides of the trimer, exposing V3 and the coreceptor binding site (12, 14, 15).

**Figure 1.**
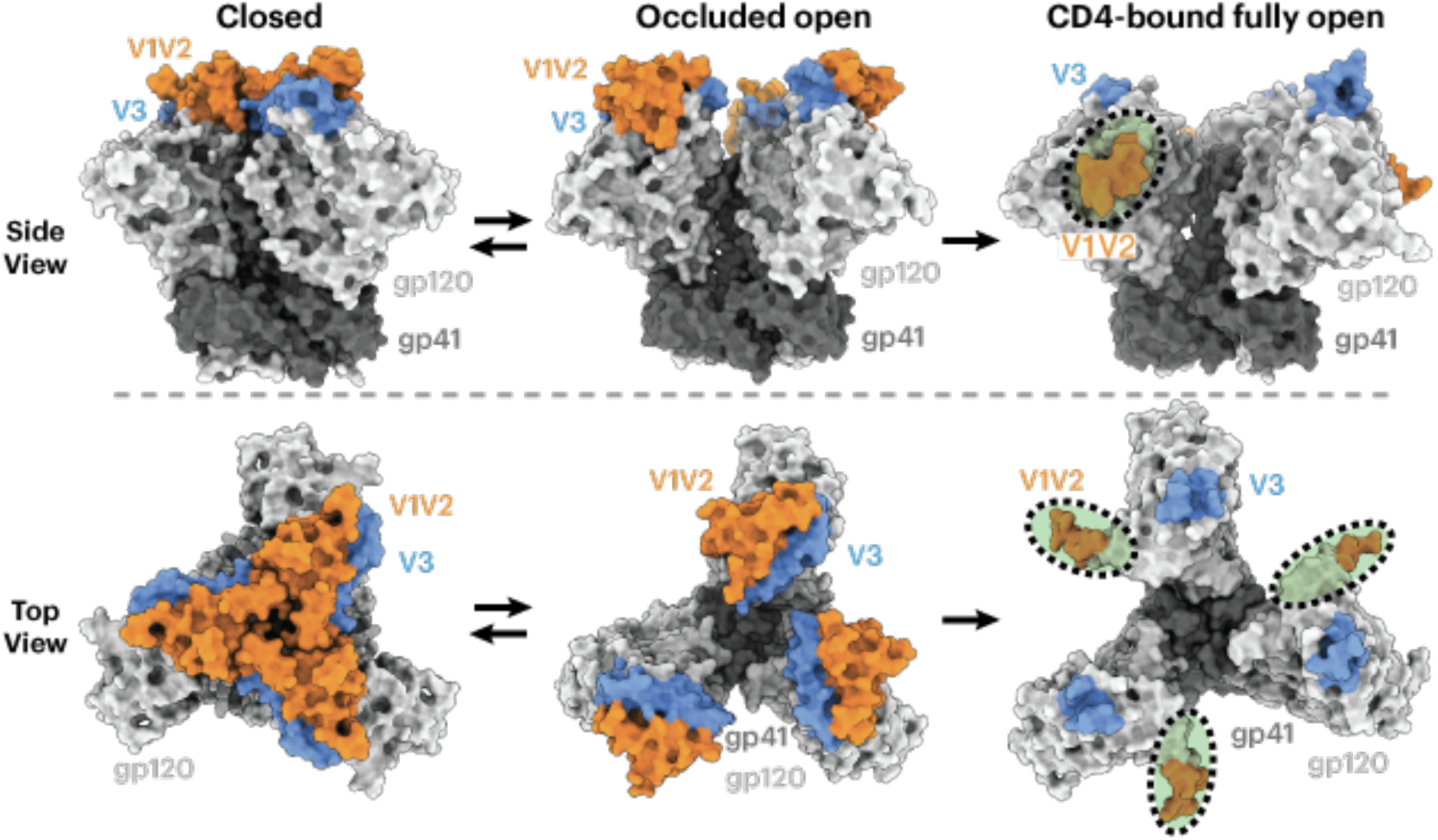
HIV Env trimers can adopt several conformational states. Env gp120 subunits are light gray, gp41 is dark gray, V1V2 regions are orange, and V3 is blue. HIV-1 Envs in different conformational states are shown in side (top) or top (bottom) views. Left: closed, prefusion Env trimer conformation with gp120 V1V2 at the apex of the trimer (PDB 5CEZ). Middle: occluded open conformation with outwardly rotated gp120 protomers but no local structural rearrangements in V1V2 or V3 (PDB 7RYU). Right: CD4-bound fully open conformation (sCD4 not shown) with V1V2 regions largely disordered and displaced from the trimer apex to the sides of the trimer and the V3 base fully exposed with the remainder of V3 disordered (PDB 6U0L). Regions that were buried in a closed Env conformation (left) but are exposed in the sCD4-bound open conformation (right) are indicated by green shading inside a dotted black oval.

HIV-1 Env is targeted by various types of antibodies, including rare broadly neutralizing antibodies (bNAbs) that neutralize multiple HIV-1 strains (16). Although the epitopes of bNAbs have been mapped to the closed, pre-fusion Env trimer (7) and an occluded-open Env conformation (12, 13), one class of antibodies, CD4-induced (CD4i) antibodies, recognize Env regions that are exposed as a result of Env conformational changes induced by CD4 binding (17). CD4i antibodies are not very potent neutralizers, perhaps because their epitope is hidden on the ligand-free, closed conformation trimer and/or because of the limited steric accessibility of the epitope when the viral Env is attached to CD4 on the host cell (18). However, because they bind to relatively conserved regions of gp120, these antibodies tend to recognize multiple HIV-1 strains (19–23).

CD4i antibodies were initially structurally characterized as complexes of an antibody Fab bound to a monomeric gp120 core that contained truncations in the N- and C-termini, V1V2, and V3, as was first demonstrated in the structure of gp120 complexed with the CD4i antibody 17b and sCD4 (24). The CD4i epitope on a monomeric gp120 core is located near the base of V3 and the gp120 bridging sheet, a four-stranded anti-parallel β-sheet comprising the gp120 β20 and β21 strands and the β2 and β3 strands at the base of V1V2. The anti-parallel four-stranded bridging sheet (β20-β21-β2-β3) configuration was later observed in sCD4-bound, open conformation SOSIP Env trimer structures (12, 14, 15, 25). However, the first closed conformation SOSIP Env trimer structures showed a rearranged three-stranded bridging sheet in which the β2 and β3 gp120 strands switched positions from the four-stranded bridging sheet conformation to a three-stranded conformation in which β21 is parallel to the β3 strand and β2 adopts a helical conformation on the opposite side of β3 (9, 10). The three-stranded sheet conformation has subsequently been observed in Env trimer structures lacking bound sCD4 (7). Although an Env trimer–sCD4–coreceptor structure is not yet available, CD4i antibodies mimic host coreceptors in that they require conformational changes within Env for binding. Some CD4i antibodies, e.g., E51 and 412d, mimic the N-terminal residues of the CCR5 coreceptor by including sulfotyrosines in their heavy chain complementarity determining region 3 (CDRH3) regions (15, 26, 27).

The gp120 V1V2 region induces antibodies in infected individuals (28), and results from the RV144 clinical vaccine trial indicated that antibody responses against the V1V2 region correlated inversely with the risk of infection (29, 30). Multiple V1V2 antibodies have been isolated from HIV-1 donors (31). Structures of Env complexed with V1V2 bNAbs, e.g., CAP256-VRC26.25 (32, 33) and PG9 (34, 35), showed a single Fab binding to the apex of a closed Env trimer. Another type of antibody that recognizes the V1V2 region, V2i antibodies, e.g., 697D, 1361, 1393A, and 830A, were isolated from human donors and recognize conformational epitopes on gp120 but not V2 peptides (36–40). These Abs are primarily derived from the VH1-69 gene segment and exhibit weak cross-neutralizing activity against neutralization-sensitive pseudotyped viruses (36–40). Given that the monomeric gp120 core in gp120-CD4i antibody complexes adopts the bridging sheet conformation found in the gp120s of open conformation sCD4-Env trimer-CD4i antibody complexes (24), we investigated whether V2i monoclonal antibodies (mAbs) target the Env V1V2 region similarly to CD4i antibodies that recognize open conformation Envs. Here, we present single-particle cryo-EM structures of V2i mAbs in complex with a sCD4-bound open SOSIP Env. For comparison, we also structurally characterized CG10, a CD4i antibody isolated from a gp120-sCD4 immunized mouse that was developed to characterize conformational rearrangements of gp120 associated with receptor binding (41, 42). Here it is demonstrated that CG10 can directly engage both gp120 and CD4 in an open Env trimer-sCD4 complex. The structures demonstrate a variety of binding poses for these antibodies, all of which can be classified within the CD4i class of antibodies that recognize a sCD4-bound open Env trimer conformation.

## Results

### Binding experiments demonstrate that V2i mAbs require sCD4 for recognizing Env trimer

To investigate whether V2i mAbs recognize closed or sCD4-bound open Envs, we evaluated binding of V2i antibody Fabs to either a ligand-free, closed B41 SOSIP Env trimer or to a sCD4-bound open trimer. C-terminally tagged soluble SOSIP trimers were immobilized on an ELISA plate using an anti-tag antibody, and the binding of a V2i Fab to trimer was subsequently detected in the presence or absence of sCD4 (Fig. 2A). As a positive control for CD4i antibody binding, we used a Fab from 17b, whose epitope is buried in a closed Env trimer but exposed in an open trimer and on gp120 monomers (12, 14, 24). As expected, 17b showed enhanced binding in the presence of sCD4 (Fig. 2B). The same behavior was observed for Fabs from V2i mAbs 1361, 1393A, 830A, and 697D (36–40); thus, maximal binding was observed in the presence of sCD4. These observations suggest that these V2i mAbs recognize the Env trimer similarly to CD4i antibodies, whereas anti-V1V2 bNAbs such as CAP256-VRC26.25 or PG9 recognize the closed Env conformation (32–35). By contrast to the other V2i antibodies, 830A exhibited weak binding to the sCD4-free, closed conformation trimer (Fig. 2C). These data provide a potential explanation for a macaque simian-HIV (SHIV) challenge study showing that passively administered antibody 830A reduced viral titers (43, 44).

**Figure 2.**
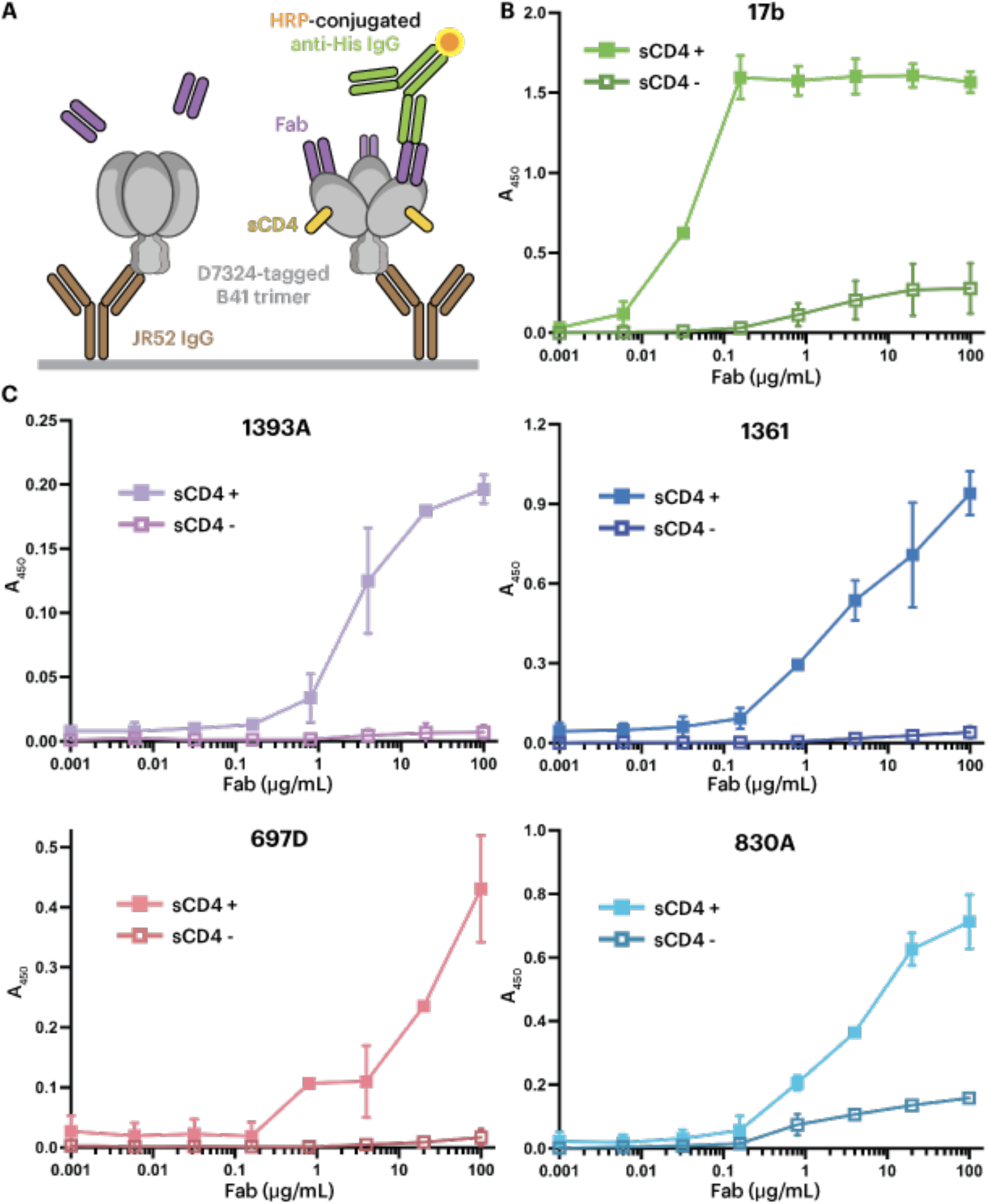
ELISAs demonstrate that V2i mAb Fabs preferentially recognize the sCD4-induced open conformation of B41 Env. (A) Schematic view of an ELISA in which D7324-tagged B41 SOSIP trimer was captured by the anti-D7324 antibody JR-52 (6), and V2i Fabs were added in the absence (sCD4-, left) or presence (sCD4+, right) of sCD4. (B) Positive control demonstrating preferential binding of the CD4i 17b Fab in the presence of sCD4. (C) Binding of the indicated V2i Fabs to B41 Env in the presence and absence of sCD4.

### V2i mAbs bind to the CD4-induced open Env conformation

To further investigate the recognition mechanism of V2i mAbs, we structurally characterized V2i Fabs complexed with a sCD4-bound BG505 SOSIP.664 Env trimer (6), an Env trimer for which structures are available in both closed and sCD4-bound open conformational states (12, 14, 15). We prepared complexes for single-particle cryo-EM by incubating a V2i Fab with BG505 SOSIP trimer and sCD4. Structures of V2i Fab-BG505-sCD4 complexes were reconstructed to 7.5Å, 6.1Å, 7.0Å, and 7.3Å overall resolutions for Fabs 1393A, 1361,697D, and 830A, respectively (Fig. S1, Table S1). Although these resolutions prohibited analyses of detailed sidechain interactions, we could identify the polypeptide backbones and improve the quality of map densities at the Fab-gp120 interaction regions using local refinement methods (45, 46). To generate coordinates, we fitted the structure of an asymmetrically open BG505 Env trimer bound to sCD4 (PDB 6U0L (15)) into the EM maps. To generate Fab coordinates, we used a crystal structure of 830A Fab (PDB 4YWG) as a template (47), aligned its sequence to those of other V2i Fabs, replaced sidechains that differed using polyalanine, and subsequently fit the Fab models into their respective EM maps.

Density maps of V2i Fab-BG505-sCD4 complexes showed that the Envs in all four structures adopted a fully-open sCD4-bound Env conformation (12, 14, 15) in which the three gp120 protomers were rotated and displaced from the trimer three-fold axis to expose the central region of Env, and the V1V2 regions were displaced from the trimer apex to the sides of the Env to expose a largely disordered V3 region (Fig. 3). In all four V2i Fab-Env-sCD4 structures, the V2i Fab contacted the V1V2 region as well as the anti-parallel β2 and β3 strands of the four-stranded bridging sheet (Fig. 3, 4). Since these regions are not accessible in the closed prefusion or occluded-open Env conformations (Fig. 1), these V2i antibodies are predicted to recognized only the open, sCD4-bound conformation of Env.

**Figure 3.**
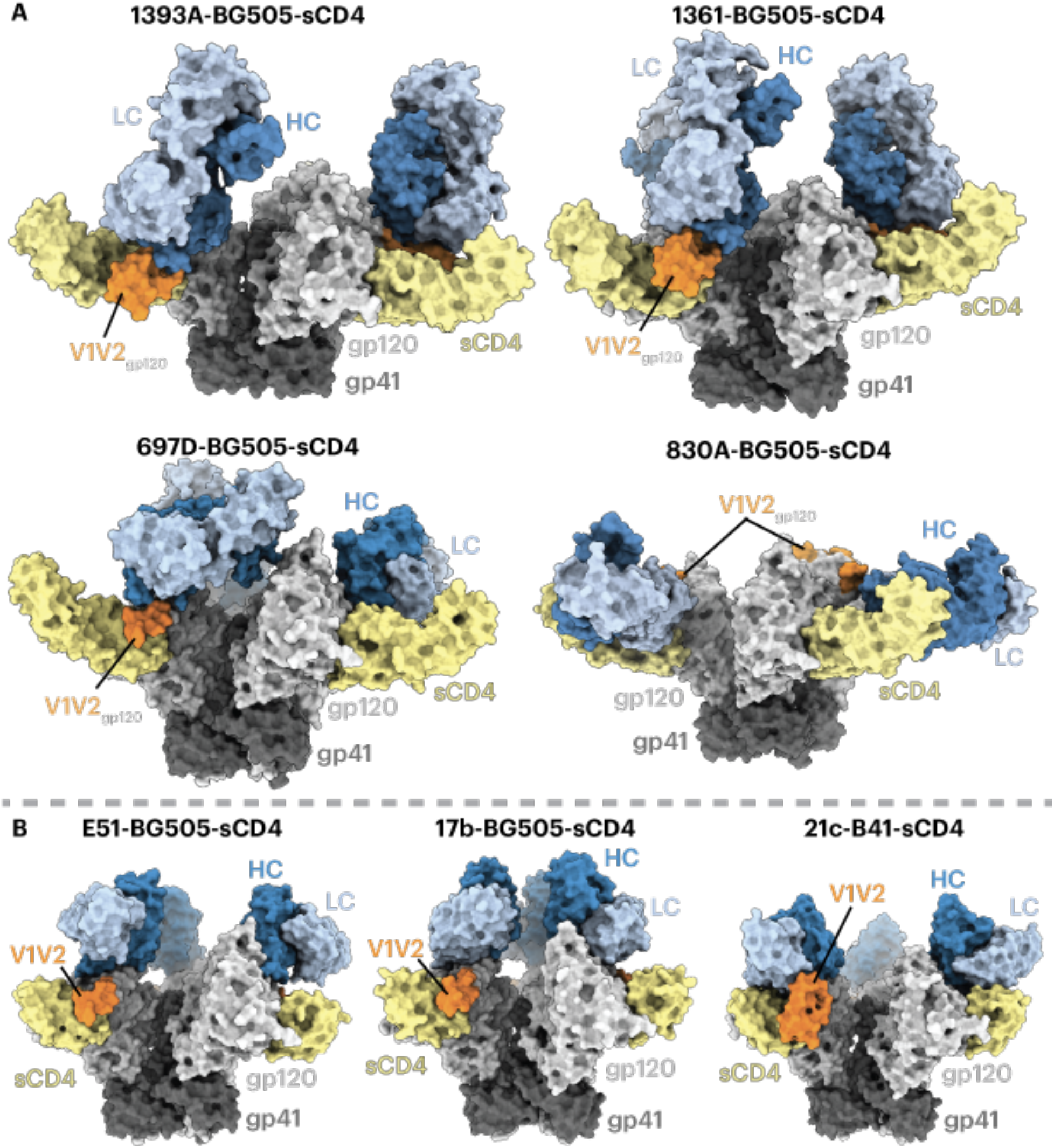
V2i Fabs adopt a variety of poses to recognize Env-sCD4. Fab HCs are dark blue, LCs are light blue, sCD4 is yellow, gp120 is light gray, gp41 is dark gray, the ordered portion of gp120 V1V2 is orange. (A) Structures are shown as surface representations. (B) Surface depictions of previous CD4i Fab-Env-sCD4 complex structures: E51-BG505-sCD4 (PDB 6U0L), 17b-BG505-sCD4-8ANC195 (PDB 6CM3), and 21c-B41-sCD4-8ANC195 (PDB 6EDU).

Antibodies 1393A and 1361 approached the gp120 β2/β3 strands and the V1V2 regions nearly vertically from the apex of an open trimer, with their V_H_-V_L_ domains facing downward (Fig. 3A, top panels). The angles of approach for 1393A and 1361 were comparable to those of CD4i Fabs such as 17b and E51, although the latter target the Env coreceptor binding site instead of its V1V2 region (12, 14, 15). Antibody 830A recognized a similar gp120 region, yet approached the sides of the gp120 protomers, resulting in a nearly horizontal alignment of the antibody V_H_-V_L_ and C_H_-C_L_ domains (Fig. 3A, bottom), whereas antibody 697D approached the V1V2 region at an angle between that seen for the 1393A and 1361 antibodies and for antibody 830A (Fig. 3A, bottom). Finally, a portion of the V1V2 region (residues Gln130_gp120_ to Glu190_gp120_) that was disordered in structures of BG505 SOSIP or B41 SOSIP Env trimers bound to sCD4 and CD4i Fabs (e.g., 17b-B41-sCD4 (PDB 5VN3) (12), 17b-BG505-sCD4-8ANC195 (PDB 6CM3) (14), E51-BG505-sCD4 (6U0L) (15)) (Fig. 3B) was resolved in the structures involving the 1393A and 1361 Fabs (Fig. 3A). An ordered portion of V1V2 was also observed in the 21c-B41-sCD4-8ANC195 structure (PDB 6EDU) in which the B41 V1V2 was stabilized by the 21c antibody light chain (14). For the 1393A-and 1361-Env complexes, the ordered portion of V1V2 was stabilized by the Fab heavy chains (Fig. 3B, 4).

**Figure 4.**
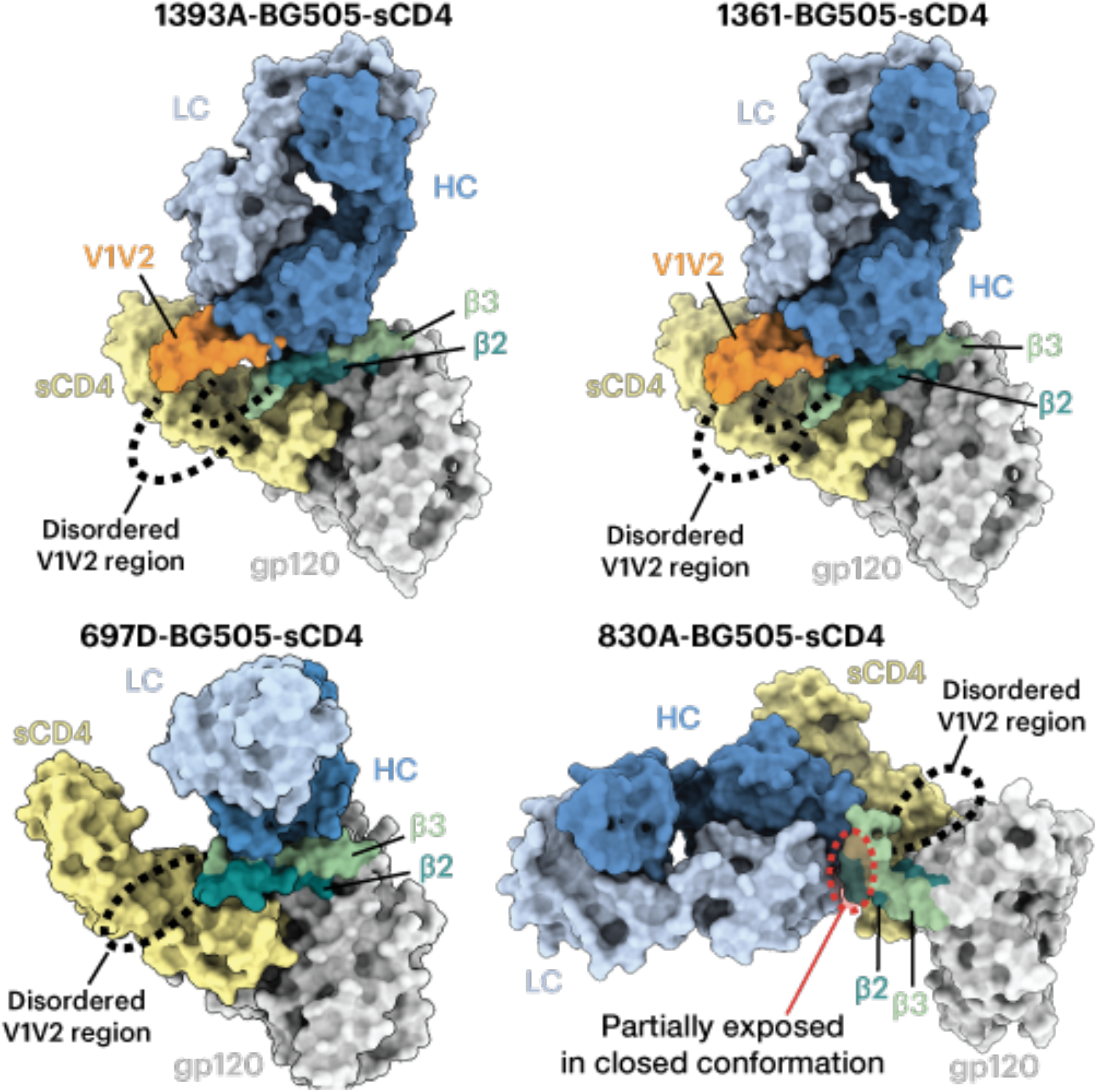
V2i Fabs contact gp120 β2, β3, and V1V2. Surface depictions of a gp120 monomer from the indicated Fab-trimeric Env-sCD4 structure are shown with the β2, β3, and V1V2 regions highlighted. Fab HCs are dark blue, LCs are light blue, sCD4 is yellow, gp120 is light gray, gp41 is dark gray, gp120 V1V2 is orange, the gp120 β2 and β3 strands are teal and green, respectively, and dashed lines indicate connections in V1V2 where density is missing in the EM maps. For the complexes with 1393A and 1361, a portion of V1V2 (orange) was stabilized by the Fab HC. Bottom right: for the complex with V2i Fab 830A, the β2 and β3 strands were displaced and bent upwards, and the 830A epitope that is partially exposed in the closed Env conformation is highlighted in a red dashed oval.

Analysis of the structure of the 830A-BG505-sCD4 complex showed that the antibody binds to a side of the V1V2 region that could be partially accessed in a closed Env trimer (Fig. 4), rationalizing the ELISA results in which weak binding of 830A to a sCD4-free, closed conformation trimer was detected (Fig. 2C).

### Comparison of Env trimer recognition by V2i antibodies and a stringent CD4i antibody

V2i antibodies elicited during natural HIV-1 infection that exhibit CD4i antibody characteristics were compared to CG10, a CD4i antibody that was elicited in a mouse immunized with a gp120-sCD4 complex (41). Unlike most other CD4i antibodies, CG10 was hypothesized to be strictly dependent on a CD4-induced conformation of gp120, typically the result of sCD4 binding (41, 42), as seen for prototype CD4i antibodies, such as 17b, that require sCD4 for binding to cell surface HIV-1 Env trimers (19) or to soluble BG505 SOSIP Env trimers (6). The requirement of sCD4 for 17b binding to Env is understood to reflect CD4-induced conformational changes, as the 17b Fab showed no direct contacts with sCD4 in crystal structures of 17b-gp120-sCD4 complexes or 17b-Env-sCD4 cryo-EM structures (12, 14, 24). By contrast, 21c, although described as a CD4-relaxed CD4i antibody that bound monomeric gp120s in the absence of sCD4, some(14), but not all (48), studies, showed contacts with sCD4 as well as with gp120 in structures of both sCD4-gp120 monomer and sCD4-Env trimer complexes (14, 48). To compare sCD4 requirements for V2i and CD4i antibodies that recognize Env trimers, we solved a 4.1 Å cryo-EM structure of CG10 complexed with sCD4 and the clade B B41 SOSIP (Fig. 5A, S2) and a 1.4 Å crystal structure of CG10 Fab (Fig. 5B, Table S2) that was fit into the CG10-B41-sCD4 density (Fig. 5A, Table S3).

**Figure 5.**
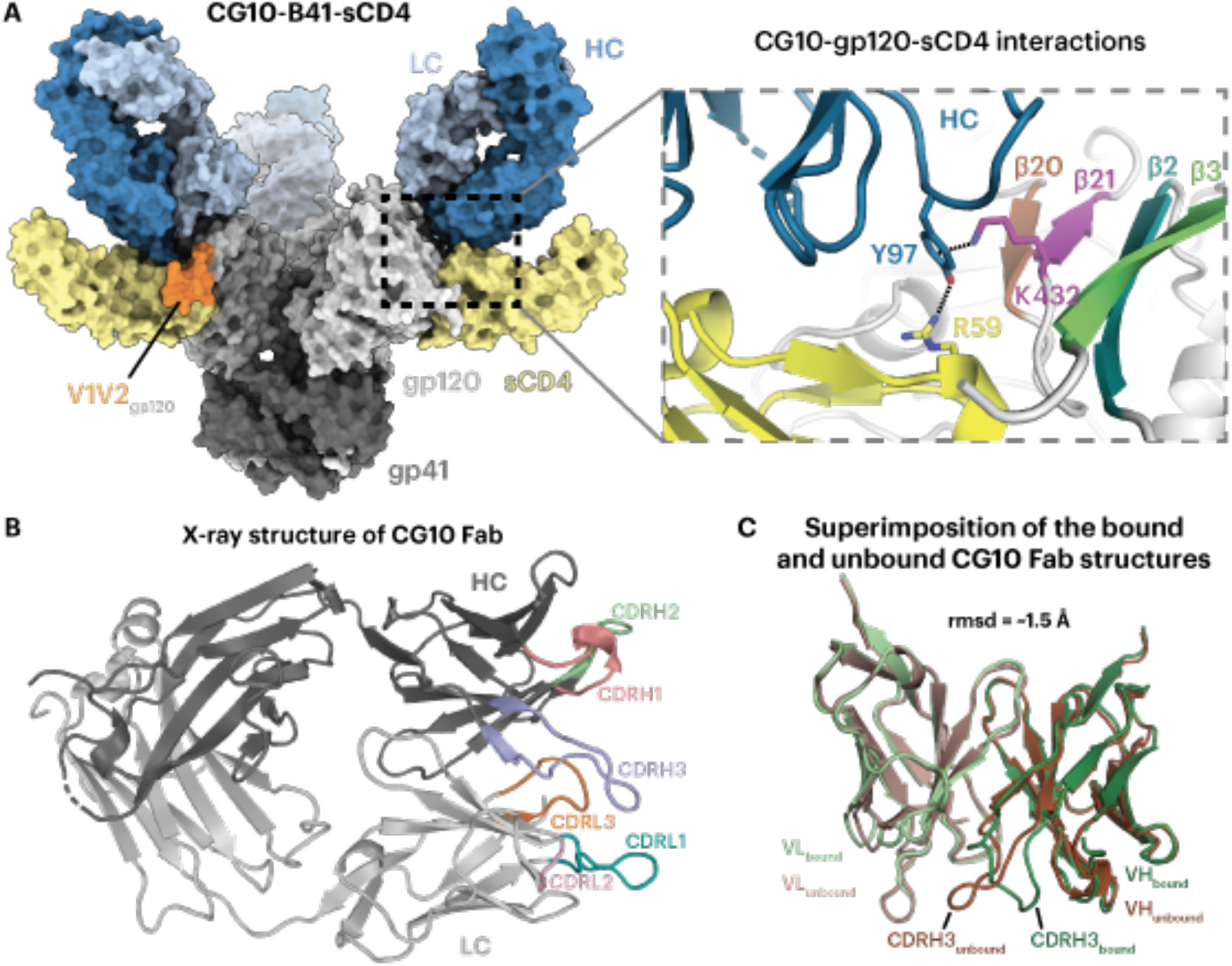
CG10 recognizes the CD4-induced open Env conformation and interacts with both sCD4 and gp120. (A) Left: surface representation of the cryo-EM structure of the CG10-B41-sCD4 complex. Env gp120 protomers are light gray, gp41 is dark gray, sCD4 is yellow, CG10 Fab HC and LC are dark and light blue, respectively, and V1V2 regions are orange. Right: Closeup of CG10 residue Tyr97_HC_ forming an interaction network with Lys432_gp120_ and Arg59_CD4_. The gp120 four-stranded bridging sheet is colored in brown, purple, teal, and green for the β20, β21, β2, and β3 strands, respectively. (B) X-ray structure of the CG10 Fab with colored CDR loops and HC and LC in dark and light gray, respectively. (C) Superimposition of the V_H_-V_L_ domains of the unbound (dark and light brown for V_H_ and V_L_, respectively) and bound (dark and light green) CG10 Fab.

The crystal structure of unbound CG10 Fab showed ordered complementarity determining regions (CDRs) on both the antibody heavy and light chains. Comparison of the V_H_-V_L_ domains in the bound and unbound Fab structures showed that the conformational changes came mainly from the CDRH3 loop, with the rest of CDRs remained similar when bound to the sCD4-B41 complex (Fig. 5C) (rmsd = ~1.5 Å for superimposition of 235 V_H_-V_L_ Ca residues). The CG10-B41-sCD4 complex structure revealed an unusual interaction of the CG10 Fab CDRH3 with residues from both the B41 gp120 and sCD4 in which CDRH3 residue Tyr97_CG10 HC_ was sandwiched between positively charged residues on gp120 (Lys432_gp120_) through a cation-π interaction and sCD4 (Arg59sCD4) through a hydrogen bond (Fig. 5A). This architecture requires the existence of positively-charged residues from both gp120 and sCD4 at their respective positions, likely explaining why CG10 was classified in the “stringent” class of CD4i antibodies (42). The specificity of this distinct interaction was demonstrated by binding experiments in which CG10 bound with higher affinity to gp120s from subtype B strains such as JR-FL and YU2, compared to a gp120 from subtype A strain BG505 (42), which can be explained by a lysine to glutamine substitution at BG505 residue 432_gp120_ that would disrupt the “sandwich” structure (Fig. 5A). Finally, the structure showed limited CG10 Fab contacts with the β2/β3 V1V2 strands that are displaced by sCD4 binding (12, 14, 15) (Fig. 5A), rationalizing previous observations demonstrating CG10 binding to gp120 cores with V1V2 truncations (42, 49).

## Discussion

The RV144 HIV-1 clinical vaccine trial showed a moderate but statistically significant decrease (31.2%) in risk of infection from viral infection (50). In that study, a robust antibody response against the gp120 V1V2 region was inversely correlated with the risk of infection (29, 51, 30, 44). While vaccination trials such as RV144 using monomeric gp120 immunogens induced antibodies with antiviral activities (29, 30), the antibodies elicited against epitopes exposed on monomeric gp120, but buried on a closed conformation Env trimer, have weak neutralizing activity compared to bNAbs that target the closed prefusion Env conformation. Nonetheless, the RV144-induced antibodies that target the V1V2 Env domain displayed anti-viral activities (52, 53). How antibodies of these specificities could protect against, or mediate infection has not yet been resolved. To better understand the nature of these antibodies, we characterized four mAbs isolated from HIV-1 infected donors that target the V1V2 region of gp120 that preferentially bind gp120 monomers rather than Env trimers (36–40). These V2i antibodies displayed little or no neutralizing activity *in vitro* in the absence of sCD4 but could mediate weak but detectable neutralization in the presence of sCD4 (54). They also mediated other antiviral activities, such as blocking the binding of the α4β7 integrin (a receptor on a subset of CD4+ T cells that potentially facilitate HIV-1 seeding and replication in mucosa (55–58), virus capture (59) and Fc-dependent anti-viral effector functions (58).

Binding experiments suggested that the V2i antibodies preferentially recognize a CD4-bound open Env trimer conformation (Fig. 2C). Structural analysis using single-particle cryo-EM validated the binding results by showing that the V2i epitopes were inaccessible in the closed prefusion Env conformation (Fig. 3,4). Accordingly, these V2i antibodies can be categorized as CD4i mAbs that target the V1V2 region of sCD4-bound Env. For comparison, we also structurally characterized a CD4i antibody, CG10, that was isolated from a mouse that was immunized with a gp120-sCD4 complex for the purpose of revealing potential gp120 epitopes associated with CD4 binding (41, 42). An apparently CD4-induced epitope on gp120 monomers had been generated by gp120 binding to M2, a short peptide that bound gp120 without occluding the CD4 binding site (60). Thus, binding of CG10 to the gp120–M2 complex illustrates that CD4 itself is not absolutely required for CG10 binding to monomeric gp120. However, the structure of CG10-B41-sCD4, a complex of an Env trimer bound to both CG10 and sCD4, revealed a direct interaction between the antibody and sCD4, which further explains the reported stringency of the requirement for sCD4 for binding of CG10 to HIV-1 Env (41, 42) (Fig. 5A).

Unlike CG10, which was generated in response to recognition of an injected sCD4-gp120 complex (41), the V2i antibodies characterized in this study were isolated from HIV-1 infected donors, raising the question of how these antibodies were elicited since their epitopes are not completely accessible on closed, prefusion Env trimers (Fig. 1). Here, we propose two scenarios that could lead to the production of this type of antibody:

First, although cryo-electron tomography studies of Env conformations on HIV-1 virions show Env trimers in a closed, prefusion conformation (61–64), it is possible that ligand-free Env trimers can transiently adopt an open conformation that mimics a CD4-induced open conformation. This hypothesis is supported by studies of the dynamics of Env trimers showing structural changes occurring in the absence of CD4 that suggest that trimers can be in equilibrium between closed and more open conformations (65). Support for this model comes from biological studies. For example, studies focused on the Env V1V2 domain showed that the relative levels of V2i mAb binding to Env-transfected cells increases with increasing time of exposure of cells to the mAb (54). The transient exposure of cryptic epitopes on unliganded trimers is further supported by data showing that, V2i antibodies displayed an increased ability to neutralize virus if exposure of virus to V2i antibodies is extended beyond the one-hour incubation time typically used in *in vitro* neutralization experiments (54); this suggests that during the extended incubation period, epitopes occluded in the closed trimer become exposed for periods long enough to be recognized and bound by antibodies. These data showing that unliganded trimers can undergo transient antigenic alterations imply that they can also present transiently exposed epitopes as immunogenic determinants that can induce an immune response.

A second possibility is that V2i and other CD4i antibodies are elicited by CD4-gp120 complexes on target cells that form after gp120 is shed during fusion between the host and viral membranes, as suggested by electron tomography imaging of virus-host cells linked by prehairpin intermediates trapped by fusion inhibitors (66) (Fig. 6). In this scenario, CD4 binding followed by coreceptor binding to Env induces trimer opening (Fig. 6A), formation of the pre-hairpin intermediate in which the Env fusion peptide is inserted into the host cell membrane (66), and shedding of gp120 protomers (Fig. 6B). This process would generate membrane-associated CD4-gp120 complexes that remain on the surface of the host cell after fusion (Fig. 6B), serving as antigens that could elicit V2i and CD4i antibodies, and could function as targets for Fc-dependent antiviral activities of V2i antibodies (58).

**Figure 6.**
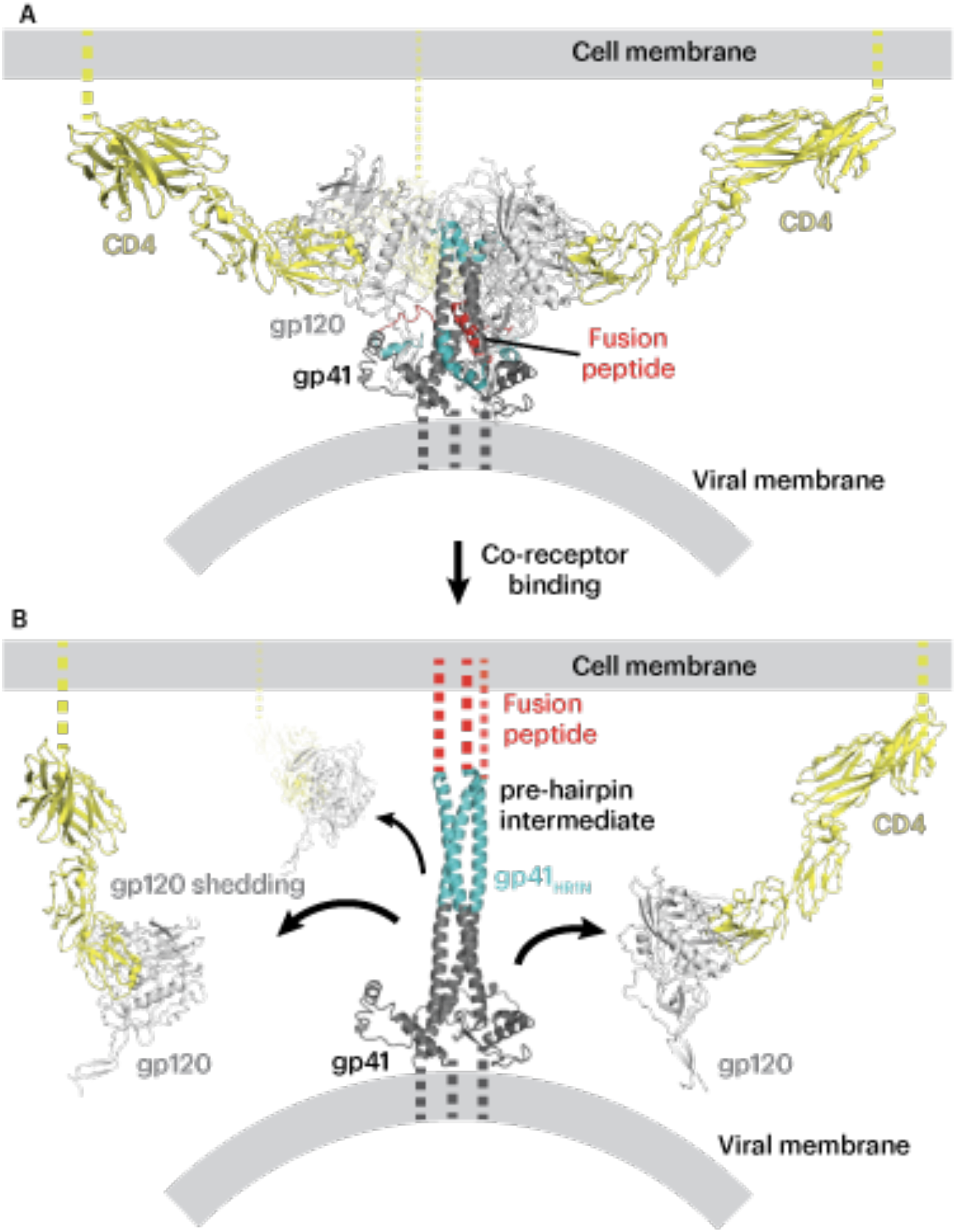
Model of post-infection formation of CD4-gp120 complexes on the host cell. (A) Open conformation Env trimer (light and dark gray with fusion peptide in red and the N-terminus of gp41 in cyan) on a viral membrane after binding to the host cell CD4 (yellow) on the host membrane. (B) A model of the pre-hairpin intermediate structure linking the viral and host cell membranes that is formed after host cell coreceptor binding (66). As potential targets to elicit V2i and CD4i antibodies during HIV-1 infection, cell surface CD4-gp120 complexes formed after gp120 shedding (black arrows) are shown on the host cell membrane.

## Methods

### Protein expression and purification

Native-like HIV-1 Env trimers SOSIP.664 (BG505 and B41-D7324) contained ‘SOS’ mutations at positions A501C_gp120_ and T605C_gp41_, the ‘IP’ mutation (I559P_gp41_), an improved furin-cleavage site (REKR to RRRRRR), and a truncation after gp41 residue 664 (6, 8). For BG505, an additional mutation was introduced to include a potential *N*-linked glycosylation site at residue 332_gp120_ (T332N_gp120_). For B41 including the D7324 tag (6), the GSAPTKAKRRVVQREKR tag sequence was added after residue 664 in the gp41 ectodomain. Both Env trimers were expressed in Expi293F cells as described (13, 14). Transfected cell supernatants were filtered and Env trimers were purified as described (13) by a 2G12 immunoaffinity column followed by size-exclusion chromatography (SEC) using a Superose 6 16/600 column (Cytiva). B41 SOSIP.664 v4.2 (8) was expressed in CHO stable cell lines kindly provided by Al Cupo and John Moore (Weill Cornell Medical College) and purified as described for BG505 SOSIP trimers produced in Expi293F cells.

All Fabs and sCD4 proteins were expressed in Expi293F cells. Fab light chain and 6x-His tagged heavy chain expression vectors were co-transfected and purified by Ni-NTA chromatography followed by a Superdex 200 Increase 10/300 GL column (Cytiva) SEC as previously described (13). The sCD4 expression vector encoded the D1D2 subunits of sCD4 (D1D2 domain residues 1-186) followed by a Strep II Tag. sCD4 protein was purified by StrepTrap HP affinity columns (Cytiva) followed by SEC using a Superdex 200 Increase 10/300 GL column (Cytiva).

### ELISA for mAb binding of CD4-induced Env

ELISAs were conducted by coating Corning Costar 96-Well Assay high binding plates (07-200-39) with the JR-52 mAb (kind gift of James Robinson, Tulane University), a mouse IgG that recognizes the D7324 tag (6), at 5 μg/ml in 0.1 M NaHCO_3_ (pH 9.6) and incubating at 4°C overnight. Excess JR-52 mAb was removed and plates were blocked for 1 hour at room temperature with 3% BSA in TBS-T (20 mM Tris, 150 mM NaCl, 0.1% Tween20). Blocking buffer was removed and D7324-tagged B41 SOSIP was added at 5 μg/ml. After a 1-hour incubation at room temperature, B41-D7324 was removed. For some experiments, sCD4 was added at 100 μg/ml and incubated for 2 hours at room temperature. His-tagged Fabs were serially diluted with 3% BSA in TBS-T at a top concentration of 100 μg/ml and incubated for 2 hours at room temperature. Fabs were removed and plates were washed twice with TBS-T. Mouse anti-His tag mAb conjugated with horseradish peroxidase (GenScript: A00186) at a 1:8000 dilution was added and incubated for 30 minutes at room temperature. Plates were then washed 3 times with TBS-T. Colorimetric detection was accomplished using 1-Step™ Ultra TMB-ELISA Substrate Solution (ThermoFisher Scientific: 34029) and color development was quenched with 1.0 N HCl. Absorption was measured at 450 nm. Two independent biological replicates for each ELISA experiment were performed.

### Cryo-EM sample preparation

V2i Fab-BG505-sCD4 and CG10-B41-sCD4 complexes were prepared by incubating purified and concentrated Fabs with soluble trimers and sCD4 at a molar ratio of (3.6:1:3.6 Fab:Env:sCD4) at room temperature for 4 hours. A final concentration of 0.02% (w/v) fluorinated octylmaltoside (Anatrace) was added to samples before cryo-freezing. Cryo-EM grids were prepared using a Mark IV Vitrobot (ThermoFisher) operated at 12°C and 100% humidity. 2.5 μL of concentrated sample was applied to 300 mesh Quantifoil R1.2/1.3 grids, incubated for 20 seconds and blotted for 4 seconds, and grids were then plunge frozen in liquid ethane that was cooled by liquid nitrogen.

### Cryo-EM data collection and processing

Cryo-grids were loaded onto a 300kV Titan Krios electron microscope (ThermoFisher) equipped with a GIF Quantum energy filter (slit width 20 eV) operating at 105,000x magnification (nominal). Defocus ranges for all complexes were set to 1.8-3.0 μm. Movies for 1393A, 1361, 697D, and 830A complexes were recorded with a 6k x 4k Gatan K3 direct electron detector operating in super-resolution mode with pixel size of 0.416 Å•pixel^-1^ using SerialEM v3.7 software (67); movies for CG10-BG505-sCD4 complex were recorded with a 4k x 4k Gatan K2 Summit direct electron detector operating in super-resolution mode with a 0.695 Å•pixel^-1^. The recorded movies were sectioned into 40 subframes with dose rate of 1.5 e^-^/Å^2^•subframe, generating a total dose of 60 e^-^•Å^-2^. A total of 3,239 (1393A-BG505-sCD4), 2,412 (1361-BG505-sCD4), 2,340 (697D-BG505- sCD4), and 1,872 (830A-BG505-sCD4), and 3,528 (CG10-B41-sCD4) movies were motion-corrected using MotionCor2 (68) with 2x binning, and CTFs of the motion-corrected micrographs were calculated using CTFFIND v4.1.14 (69). Particles automatically picked in cryoSPARC v3.2 (45) using “Blob picker” program and classified using the “2D classification” program. Good 2D classes were selected for a second iteration of reference-free 2D classification. *Ab initio* models were generated and were subsequently refined with 3D refinements in cryoSPARC v3.2 (45, 46). 3D Fourier Shell Correlation (FSC) of maps were calculated using the Remote 3DFSC Processing Server as described (70). The quality of EM map densities for Fab-gp120 interfaces were slightly improved using the cryoSPARC “Local refinement” in which particle alignments were focused on one protomer of a Fab-gp120-sCD4 complex (45, 46).

### X-ray crystallography

Crystallization screens for CG10 Fab were carried out using the sitting drop diffusion vapor diffusion method at room temperature by mixing the Fab with an equal amount of screen solution (Hampton Research) using a TTP Labtech Mosquito automatic pipetting robot. CG10 Fab crystals were obtained in 14% (w/v) PEG 4,000, 0.1M MES (pH 6.6) at room temperature. Crystals were looped and cryopreserved in liquid nitrogen.

X-ray diffraction data were collected using a Pilatus 6M detector (Dectris) at the Stanford Synchrotron Radiation Lightsource (SSRL) beamline 12-2 at a wavelength of 1.0Å. Data were indexed, integrated and scaled in XDS (71, 72) and merged with AIMLESS v0.7.4 (73). The CG10 Fab structure was determined by molecular replacement using PHASER v2.8.2 (74) using a mouse antibody with separated V_H_-V_L_ and C_H_-C_L_ domains as the search models (PDB 4CMH). Coordinates of the Fab were refined using Phenix v1.19.2 (75, 76), and iterations of manual refinement using Coot v0.9 (77) (Supplementary Table 1).

### Model building

For V2i Fab-BG505-sCD4 complexes, coordinates for BG505 Env and sCD4 were fitted into the corresponding regions of density maps using an open conformation BG505 trimer structure (PDB 6U0L) (15). Coordinates for 830A Fab were fitted using the crystal structure of a 830A-gp120 complex (PDB 4YWG) (47). For other V2i Fab coordinates (1393A, 1361, and 697D), residues that differed from 830A were replaced by polyalanines and fitted into their respective EM maps. For the CG10-B41-sCD4 cryo-EM complex, coordinates for CG10 Fab, B41 gp120 and gp41 subunits, and sCD4 were fitted into the corresponding regions of the EM density maps using the following coordinates for initial fitting: sCD4, gp120 and gp41 from an open conformation B41 SOSIP (PDB 5VN3) (12) and the unbound CG10 Fab (this study). Iterations of whole-complex refinements were carried out in Phenix (real space refine) (76) and manually done using Coot (77).

### Structural analyses

Structural figures were made using PyMOL v2.5.1 (Schrödinger, LLC) or ChimeraX v1.2.5 (78). Interacting residues between CG10 Fab and B41-sCD4 were analyzed in PDBePISA (79) using the following definitions: potential hydrogen bonds were assigned using geometric criteria of interatomic distance of <3.5Å between the donor and acceptor residues and an A-D-H angle >90°. Hydrogen atoms were added to proteins using PDB2PQR (80). The maximum distance allowed for van der Waals interaction was 4.0Å. Rmsds were calculated for Cα atoms after superimposition in PyMOL v2.5.1 (Schrödinger, LLC) of the CG10 Fab from the CG10-B41-sCD4 complex and the unbound CG10 Fab X-ray structure.

### Data availability

Cryo-EM maps generated in this study have been deposited in the Electron Microscopy Data Bank (EMDB) with accession codes EMD-27209, EMD-27210, EMD-27211, and EMD-27212 for V2i Fab complexes 1393A-BG505-sCD4, 1361-BG505-sCD4, 697D-BG505-sCD4, and 830A- BG505-sCD4, respectively. The cryo-EM map for CG10-B41-sCD4 complex was deposited in the EMDB under the accession code EMD-27208, and atomic model coordinates were deposited in the Protein Data Bank (PDB) under the accession code of 8D5C. The X-ray structure of CG10 Fab was deposited in the PDB under the code of 8D54.

## Author contributions

Z.Y., K.A.D, J.M.G., S.Z.-P., and P.J.B. designed the research. Z.Y. and K.A.D. performed experiments and analyzed results. Z.Y., K.A.D., and P.J.B. wrote the paper with input from co-authors.

## Acknowledgements

We thank Jost Vielmetter at the Beckman Institute Protein Expression Center at Caltech for protein production, John Moore (Weill Cornell Medical College) for the B41 stable cell line, and James Robinson (Tulane University) for the JR-52 mAb. Cryo-EM studies were performed in the Beckman Institute Resource Center for Transmission Electron Microscopy at Caltech with assistance from Dr. S. Chen (director). We thank the Gordon and Betty Moore and Beckman Foundations for the gifts to Caltech to support the Molecular Observatory (Dr. Jens Kaiser, director) and the Stanford Synchrotron Radiation Lightsource (SSRL) beamline staff for data collection. Use of the SSRL, SLAC National Accelerator Laboratory, is supported by the U.S. Department of Energy, Office of Science, Office of Basic Energy Sciences under contract No. DE- AC02-c76SF00515. The SSRL Structural Molecular Biology Program is supported by the DOE Office of Biological and Environmental Research, and by the National Institutes of Health, National Institute of General Medical Sciences (P41GM103393). The contents of this publication are solely the responsibility of the authors and do not necessarily represent the official views of NIGMS or NIH. This work was supported by grants from the National Institute of Allergy and Infectious Diseases (NIAID) HIVRAD P01 AI100148 (P.J.B.) and R01AI145655 (S.Z.P.), a Gates CAVD grant INV-002143 (P.J.B.), support from the Department of Medicine, Icahn School of Medicine at Mount Sinai (S.Z.P.), and the generous support of Dr. Peter Kraus (J.M.G.).

---

**Figure S1.**
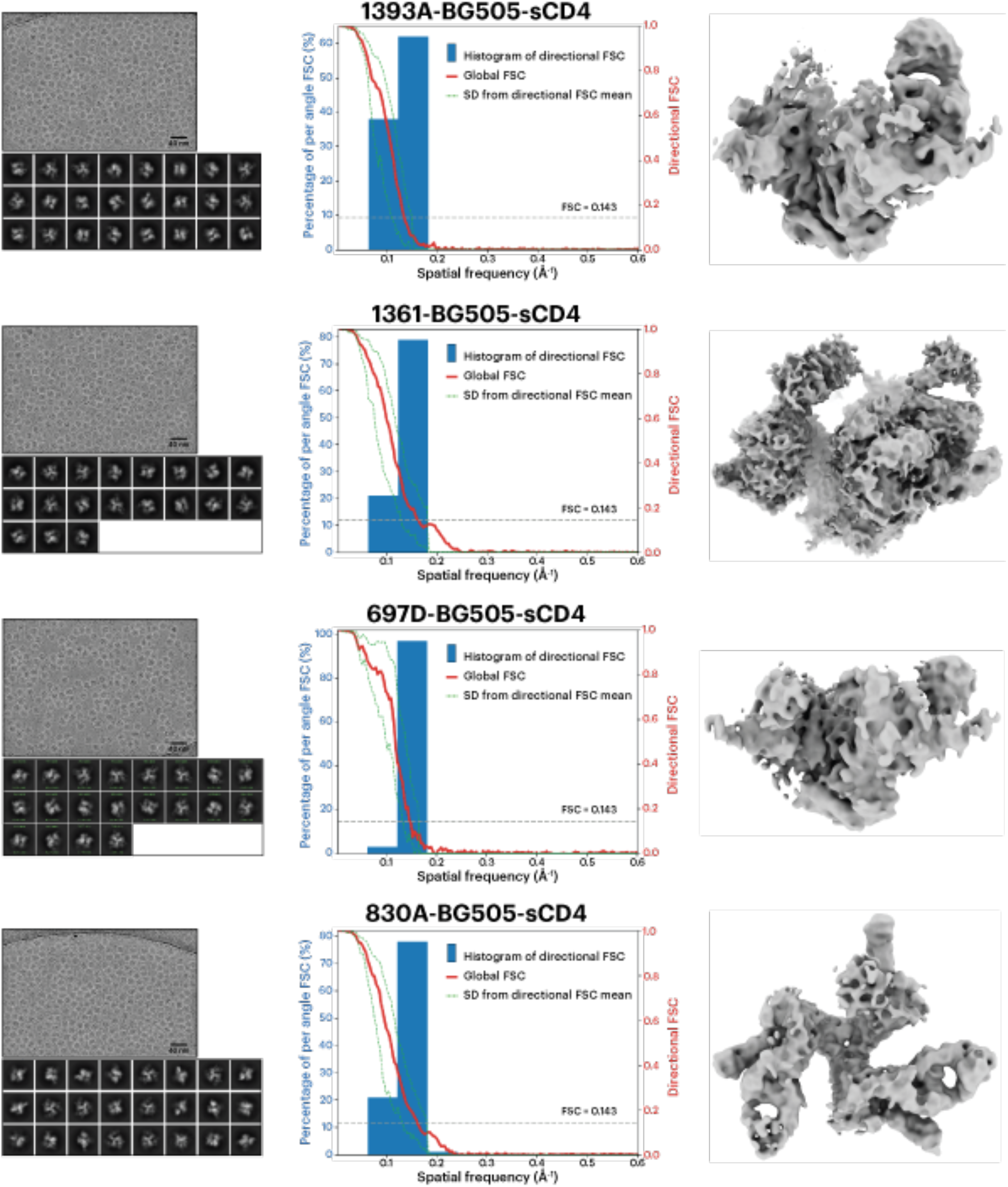
Cryo-EM data processing and validation of V2i Fab-BG505-sCD4 complexes. Left: Example micrographs and 2D class averages of V2i Fab-BG505-sCD4 complexes. Middle: Plots of global half-map FSCs (solid red line), directional resolution values ±1σ from the mean (left axis, green dashed lines) and distributions sampled over the 3D FSC (blue histograms, right axis). Right: Cryo-EM density maps of V2i Fab-BG505-sCD4 complexes.

**Figure S2.**
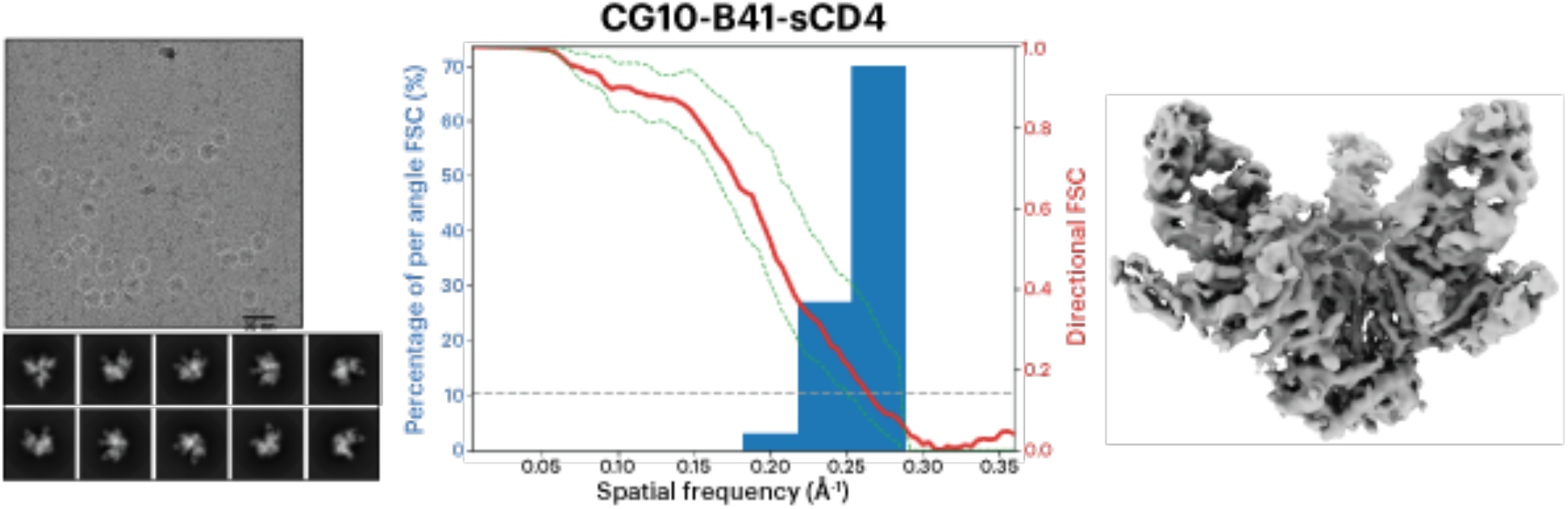
Cryo-EM data processing and validation of CG10 Fab-B41-sCD4 complex. Left: Example micrograph and 2D class averages of the CG10-B41-sCD4 complex. Middle: Plot of global half-map FSC (solid red line), directional resolution values ±1σ from the mean (left axis, green dashed lines) and distributions sampled over the 3D FSC (blue histograms, right axis). Right: Cryo-EM map of the CG10-B41-sCD4 complex.

**Supplementary Table 1.** Cryo-EM data collection, refinement, and validation statistics.

|  | **1393A-BG505-**<br>**sCD4**<br>**(EMD-27209)** | **1361-BG505-**<br>**sCD4**<br>**(EMD-27210)** | **697D-BG505-**<br>**sCD4**<br>**(EMD-27211)** | **830A-BG505-**<br>**sCD4**<br>**(EMD-27212)** |
| --- | --- | --- | --- | --- |
| **Data collection and**<br>**processing** |  |  |  |  |
| Magnification * | 105,000x | 105,000x | 105,000x | 105,000x |
| Voltage (kV) | 300 | 300 | 300 | 300 |
| Electron exposure (e-/Å<sup>2</sup>) | 60 | 60 | 60 | 60 |
| Defocus range (µm) | 1.8-3.0 | 1.8-3.0 | 1.8-3.0 | 1.8-3.0 |
| Pixel size (Å) | 0.416 | 0.416 | 0.435 | 0.416 |
| Recording mode | Super resolution | Super resolution | Super resolution | Super resolution |
| Collected movies (no.) | 3,239 | 2,412 | 2,340 | 1,872 |
| Symmetry imposed | C1 | C1 | C1 | C1 |
| Initial particle images (no.) | 943,748 | 843,403 | 367,952 | 436,527 |
| Final particle images (no.) | 107,234 | 104,002 | 21,655 | 16,575 |
| Overall map resolution (Å) | 7.5 | 6.1 | 7.0 | 7.3 |

\* Nominal magnification

**Supplementary Table 2.**
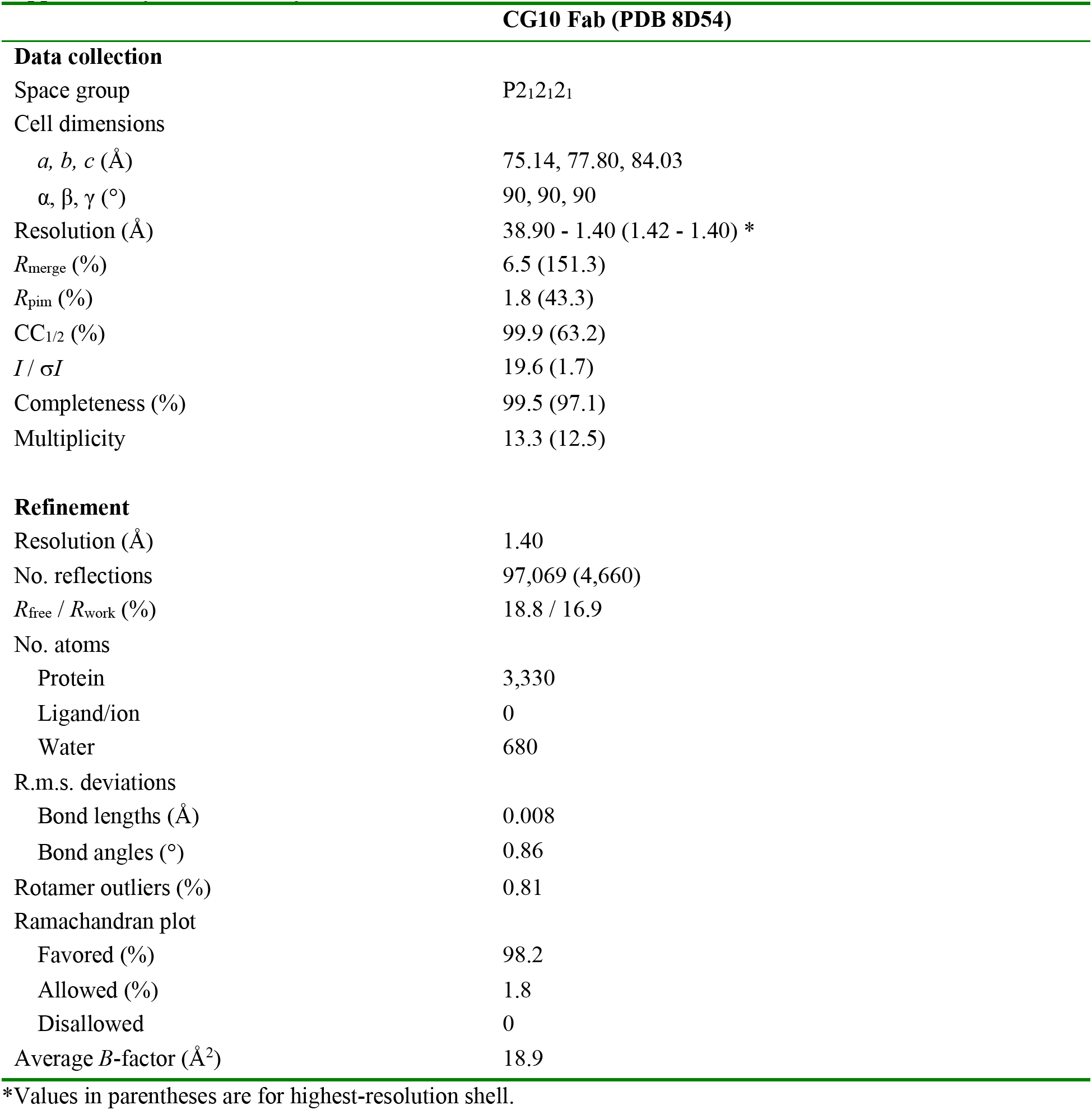
X-ray data collection and refinement statistics.

**Supplementary Table 3.**
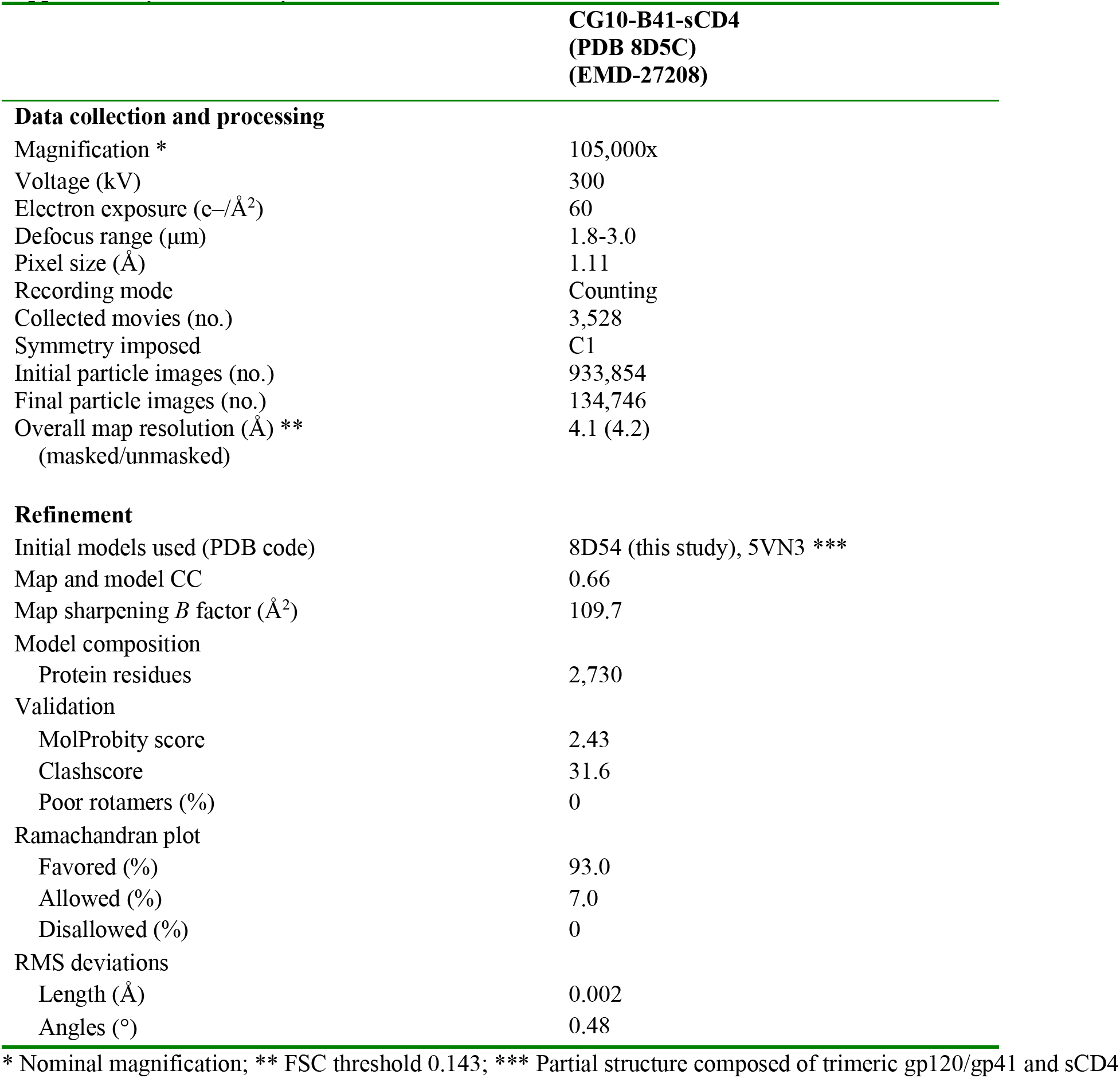
Cryo-EM data collection, refinement, and validation statistics.

